# A systematic review of sample size and power in leading neuroscience journals

**DOI:** 10.1101/217596

**Authors:** Alice Carter, Kate Tilling, Marcus R Munafò

## Abstract

Adequate sample size is key to reproducible research findings: low statistical power can increase the probability that a statistically significant result is a false positive. Journals are increasingly adopting methods to tackle issues of reproducibility, such as by introducing reporting checklists. We conducted a systematic review comparing articles submitted to *Nature Neuroscience* in the 3 months prior to checklists (n=36) that were subsequently published with articles submitted to *Nature Neuroscience* in the 3 months immediately after checklists (n=45), along with a comparison journal *Neuroscience* in this same 3-month period (n=123). We found that although the proportion of studies commenting on sample sizes increased after checklists (22% vs 53%), the proportion reporting formal power calculations decreased (14% vs 9%). Using sample size calculations for 80% power and a significance level of 5%, we found little evidence that sample sizes were adequate to achieve this level of statistical power, even for large effect sizes. Our analysis suggests that reporting checklists may not improve the use and reporting of formal power calculations.

## Introduction

The reproducibility of published scientific studies is the topic of considerable recent debate (1-5). Statistical power (the ability of a study to detect a true effect of a given magnitude) is a central feature of the null hypothesis significance testing (NHST) approach to inference that is widely used across the biomedical sciences, which if overlooked can represent a threat to reproducible research. Low statistical power of a study increases the chance that a statistically significant result represents a false positive, and that for true positives the effect size will be exaggerated from the true value (6, 7). Statistical power depends on the relationship between the sample size, the minimum effect size to be detected, and the alpha (significance) level, which is usually set to 5%.

Given the growing body of literature surrounding the impact of inadequate sample sizes and correspondingly low statistical power on scientific reproducibility, journals are becoming increasingly aware of the need to act (6, 8). One measure, taken by some journals, is to introduce a reporting checklist. This requires authors to report a range of details about their methodology and statistical analyses alongside their submission. These checklists often include a section on how sample sizes were decided on. One of the groups to adopt this policy was the Nature Publishing Group, including *Nature Neuroscience*. This was a group-wide policy that came into full effect in May 2013, although some journals, including *Nature Neuroscience*, adopted it earlier, following positive responses from a trial period (9, 10). However, these checklists may not go far enough. Goodhill (11) examined one recent issue of *Nature Neuroscience* and found that only two out of eleven studies published had either completed a power calculation or based it on pilot studies and previous studies.

We aimed to assess the impact of checklist implementation on adequate sample sizes. Articles submitted to *Nature Neuroscience* in the three-month period prior to checklist implementation were reviewed for sample sizes and power, and then compared with those submitted in the three-month period following checklist implementation. We used a negative control approach, comparing the after-checklist period in a comparator journal, *Neuroscience*, which has not implemented a checklist to date. Adequate sample size was defined by calculating the minimum sample sizes required for levels of effect sizes in different study designs, then the proportion of published studies that achieved these levels calculated.

## Methods

### Study eligibility

We focused on *in vivo* data analysis studies, including studies on human participants. A study was defined as being *in vivo* if experimentation was completed on a live subject, even if the analysis was completed on a sample taken from the subject, or if they were euthanised after the experimental condition was applied. A data analysis study was defined as a study that presented quantitative analysis of findings. Those that only presented quantitative analysis of demographic or baseline characteristics were excluded from quantitative analysis of adequate sample size, but included for qualitative analysis on sample size comments and power calculations. Functional magnetic resonance imagining (fMRI) studies were not included unless quantitative analyses were presented alongside fMRI scans. Studies that presented both *in vivo* and ex vivo or in vitro analyses were included in this review; however, only the *in vivo* results were considered for analysis.

We considered studies submitted to the journals *Nature Neuroscience* or *Neuroscience*. Studies were eligible if they had submitted to *Nature Neuroscience* in the 3 months prior to checklist implementation (October to December 2012) and those submitted in the 3 months immediately after checklist implementation (January to March 2013). Studies submitted to *Neuroscience* were eligible if they have submitted in the period from January to March 2013. The eligibility of studies is detailed in Figure 1 below, detailing at which stage studies were excluded. At each screening stage, studies could be identified as no longer eligible, as outlined in the flowchart. This was marked on the database, and a comment included as to why it had been determined as ineligible.

**Figure 1:**
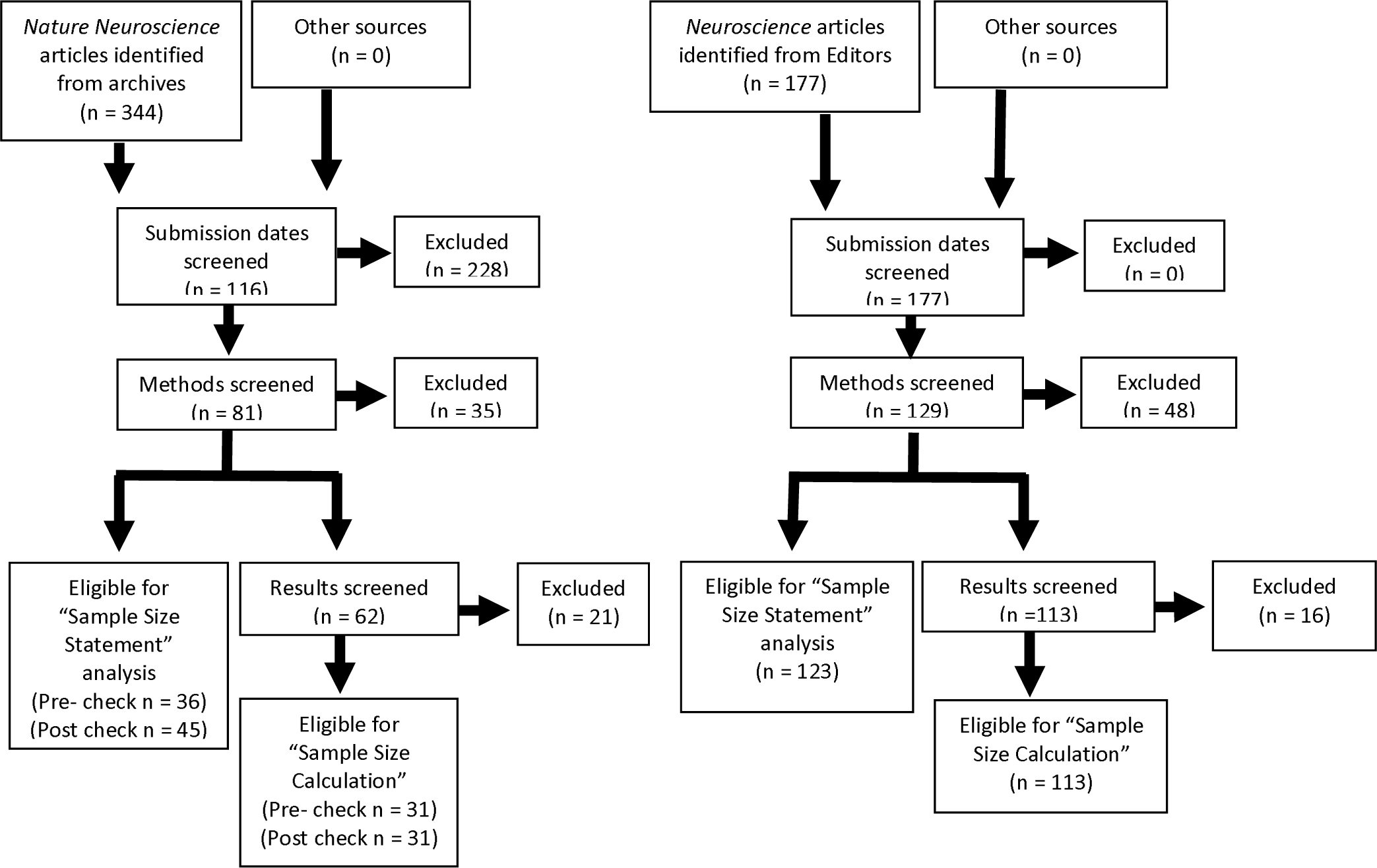
Flow chart detailing the study identification, exclusion process, review process and analysis process of submitted articles to the review and the subsequent number of articles included for each analysis.

### Information sources

All eligible studies were identified directly from the included journals. The decision to select studies based on the submission date, as opposed to the publication date, was due to the way checklists were implemented in *Nature Neuroscience*. All *Nature Neuroscience* articles publish the submission date alongside the publication date. This allowed articles to be screened directly from the *Nature Neuroscience* archives. All *Nature Neuroscience* issues from between October 2012 and March 2013 were screened for eligible studies. In the last 3 issues (January-March 2014) no eligible studies were identified, so it was deemed appropriate to end screening at this point. Articles published by *Neuroscience* do not publicly release the submission date for each article. The editors of *Neuroscience* were approached, and a full spreadsheet of all articles submitted between October 2012 and March 2013 was provided. These articles were then screened for eligibility.

### Study selection and data collection process

All articles identified that were submitted within the dates of interest were entered to a database, with key information collected, including details on the title, authors, date of submission, date of publication and journal submitted to.

Articles were initially screened to assess whether they met the primary eligibility criteria of being an *in vivo* data analysis study. Those that were eligible were screened for a second stage, where the abstract, methods and results (including tables and figures) were fully reviewed, to ensure all other eligibility criteria were met. All results presented in the main articles were extracted. At this stage, the data extracted included: sample sizes, effect estimates, and P values for each individual result presented. Only results in the main paper were assessed, and not those in supplementary materials.

Next, articles were further screened to extract data on the completion of sample size or power calculations. Where these were not presented, any comment reported on the sample size used or the subsequent power of the study was extracted, or simply reported as not making any comment. The data extracted from fully eligible studies included: the largest sample size reported of any analysis, any comment on sample size in any part of the main paper, sample size or power calculation details where they were provided, the study species and populations where human participants were involved, any comments on excluded data, ethical statements, investigator blinding, and any other comments of interest.

For quantitative analysis, and to determine whether a study had adequate power, the largest reported sample size was used. The largest sample size was determined in a number of ways due to inconsistencies in reporting and different experimental methods being used. The hierarchical process to determine this is outlined in Box 1.

#### Box 1: Systematic determination of the largest reported sample sizes across studies included in eligible studies.

1. Largest number of animals reported for experimentation:

1.1. Either from experimental or control group, since it is appropriate to maximise the sample size of the control group to increase study power.
1.2. The largest single sample size is chosen because of the way power calculations are being determined, even if two groups being compared would equal a larger total for group1 + group 2 for the analysis in the paper.
2. Where number of cells/slices are reported in conjunction with sample sizes:

2.1. Largest sample size of whole animals.

2.1.1.e.g. “8 animals were used in analysis” is selected over “10 slices from 6 animals”
2.2. If all are cells/sections of animals, the largest number of animals the samples are taken from.

2.2.1.e.g. “10 slices from 6 animals” is selected over “20 slices from 2 animals”
2.3. If all cells/sections of animals and sample from same number of animals: largest number of samples from the largest number of whole animals

2.3.1.e.g. “10 slices from 6 animals” is chosen over “10 slices from 2 animals”
3. Where no whole animal numbers are reported, and sample size is only presented as a range:

3.1. The largest number of the range is selected

3.1.1.A range of 9-11 animals, 11 is determined as the sample size

The database was then converted to a Stata file and statistical analysis was carried out using the Stata 14 statistical software package.

### Risk of bias in individual studies and across studies

Given that this review was based around reporting of results, all studies were included regardless of any potential biases. Due to poor reporting in many cases, as this review highlights, data were often hard to identify and some extracted data may have inaccuracies. We anticipated reporting bias to be present in these studies, where large effect sizes with small P values are presented, along with selective reporting of positive results. However, given that identifying these issues within the data was of principle concern in this review, all studies have been included and no further explorations of these biases has been carried out.

### Summary measures

Initially, we were Interested in all summary measures reported. However, for the analysis in this review, we restricted our measure of interest to the largest reported sample size of any analysis within each study.

## Results

### Study characteristics

A total of 204 studies were included in the final review. The majority of studies were on rodents, primarily rats and mice (69%), followed by studies on humans (18%), other mammals (8%), and other animals (5%). Of the studies eligible for qualitative analysis, 123 (60%) came from the journal *Neuroscience*, compared with 36 studies (18%) submitted before the use of checklists and 45 studies (23%) submitted after checklists were introduced in *Nature Neuroscience*. This is attributable to the higher publication frequency of *Neuroscience* compared with *Nature Neuroscience*. When considering the eligible studies for quantitative analysis, 113 studies (65%) published in *Neuroscience* were eligible, compared with 31 studies (18%) for each of the groups in *Nature Neuroscience*.

### Justification of sample size and power calculations

Across all journals, 65 studies (32%) commented on the sample size of their study. Figure 2 shows that following checklist implementation, a greater proportion of *Nature Neuroscience* studies made comments on their sample sizes, an increase from 22% to 53%. Overall, sample size was also addressed more in *Nature Neuroscience* articles than *Neuroscience*, indicating the potential influence the publishing journal can have on reporting behaviour. However, we observed a decrease in the number of formal power calculations from 16% to 9%, following the implementation of checklists.

**Figure 2:**
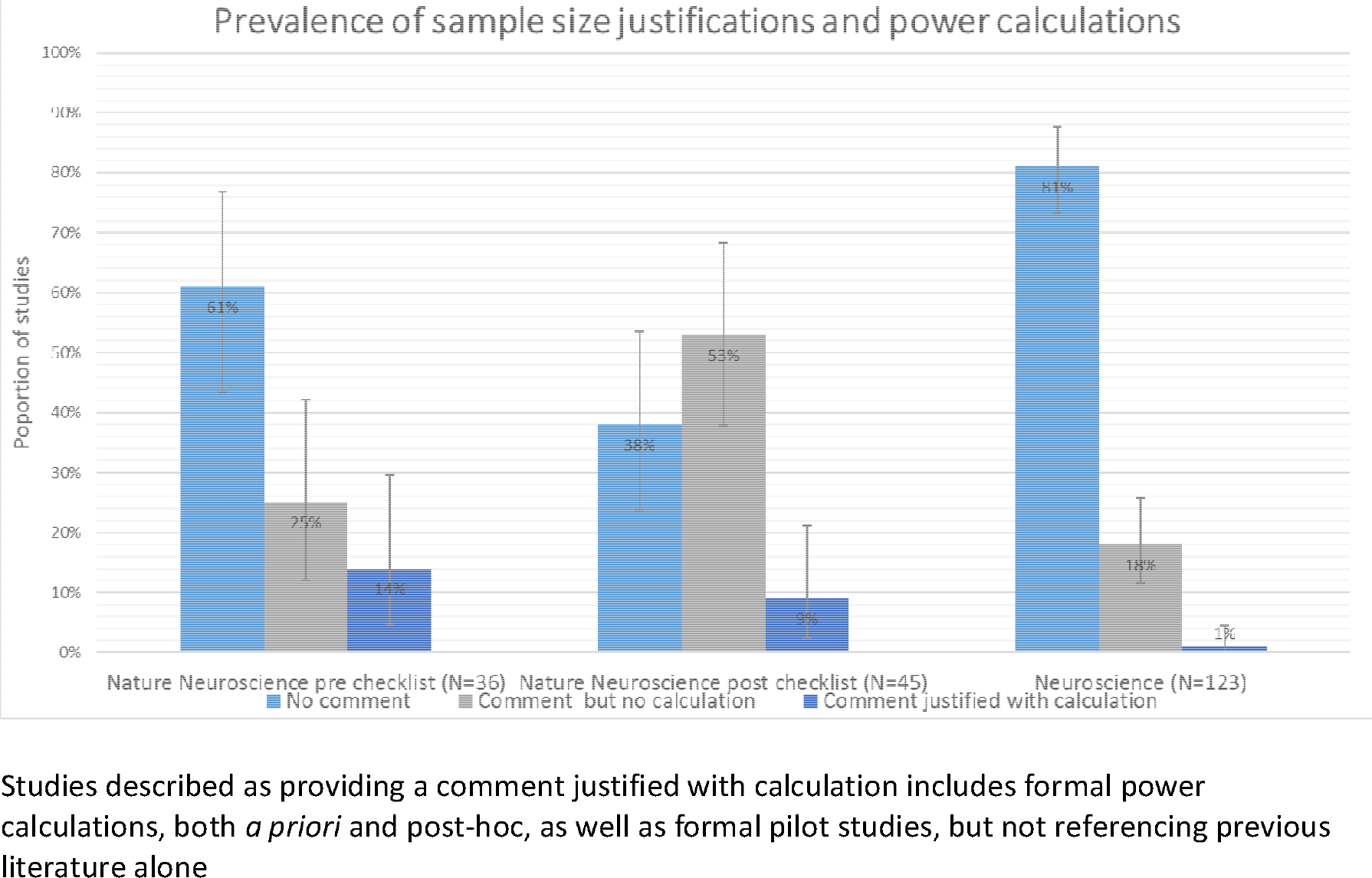
Bar chart showing prevalence of reporting behaviours on sample sizes and power calculations in the main text of eligible articles in each reviewed journal. Studies described as providing a comment justified with calculation includes formal power calculations, both a priori and post-hoc, as well as formal pilot studies, but not referencing previous literature alone

Of the 24 studies in *Nature Neuroscience* with a sample size comment after checklists were implemented, 16 studies (67%) used the words “no statistical methods/test were used”, whilst one specified that no power calculation was carried out. Eighteen (74%) of the 24 studies commented that their sample size was similar to the field or based their justification on previous studies. One study justified their sample size because they found statistically significant results: “This sample size was considered as sufficiently large because … Figure 1 B-D was significantly reduced at N = 10 flies P ≤ 0.0001”. Finally, one study determined their sample size was appropriate because “attention was paid to use only the number of mice requested and necessary to generate reproducible and reliable results”. Four studies (9%) in this group statistically justified their sample sizes; 3 studies carried out power calculations, whilst 1 carried out pilot studies to determine their sample size. Seventeen studies (38%) made no reference to their sample size.

Of those studies published before checklists in *Nature Neuroscience*, 22 studies (61%) made no comment on how their sample size was determined. Of the 9 studies that did not justify their sample size by means of a power calculation, either a priori or post hoc, or via pilot studies, 7 studies (19%) based their sample size on previous studies. Five studies (14%) in this group used power calculations to justify their sample sizes.

The ethical implications of sample size were a common theme in *Neuroscience* articles. Aside from the one study that conducted a power analysis, all other studies that justified their sample size (22 studies, 18%) referred to the need to minimize the number of animals used from an ethical point of view. Of these, 2 studies commented that they minimized the number of animals necessary to maintain statistical power. However, the remaining studies did not address issues with power and minimizing sample sizes. A total of 100 studies (81%) made no reference to justifying their sample sizes.

### Sample size calculations for adequate power

Figure 3 shows that most studies reviewed had an adequate sample size to detect only large differences for each statistical test. As anticipated, very few studies had an adequate sample size to detect small effect estimates for either test. There was no clear evidence of increased sample sizes in *Nature Neuroscience* following checklist implementation. For almost all potential effect sizes, and for all methods of analysis, there was no discernible difference in the proportion of studies powered to detect the effect size before and after the checklist.

**Figure 3:**
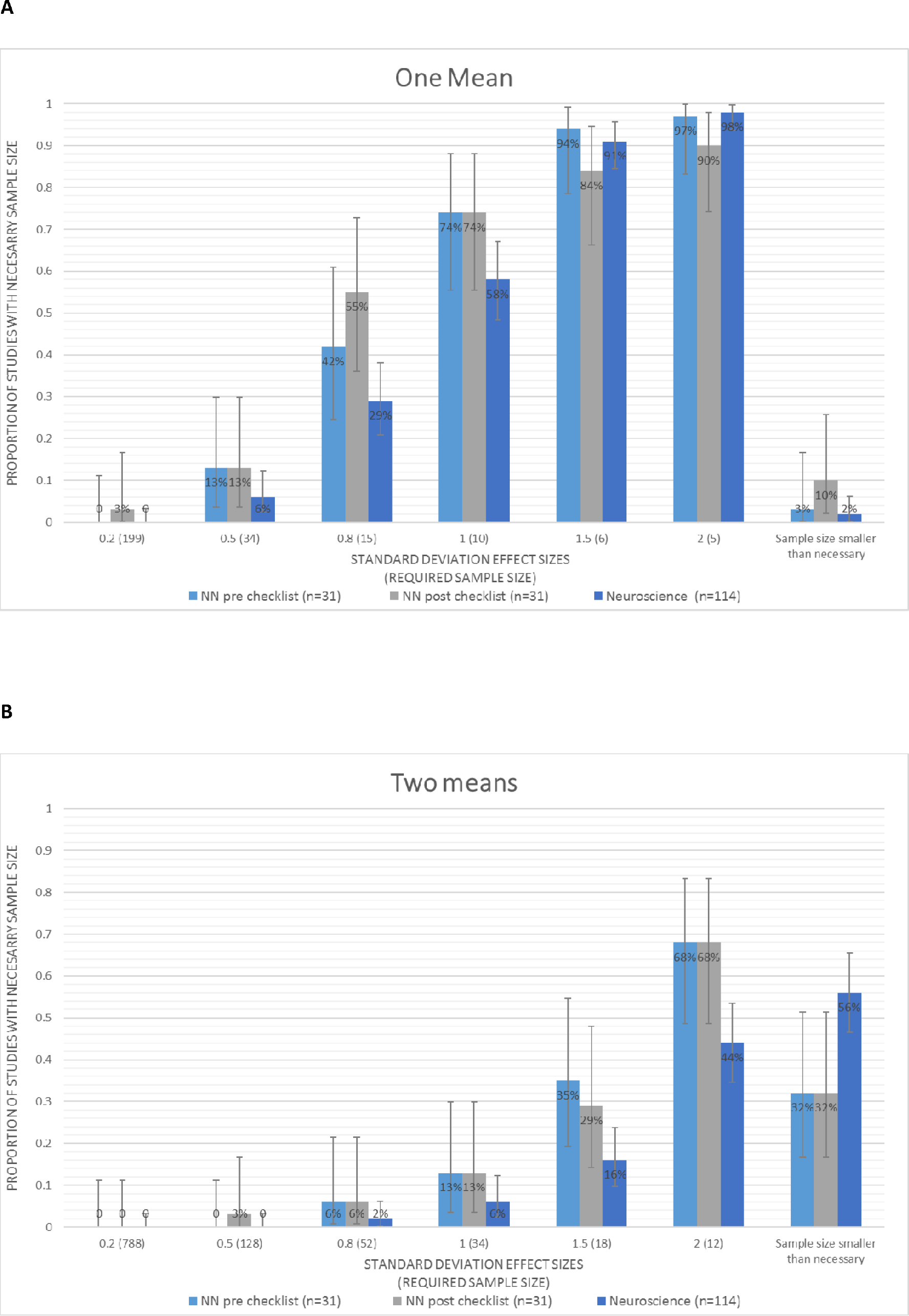
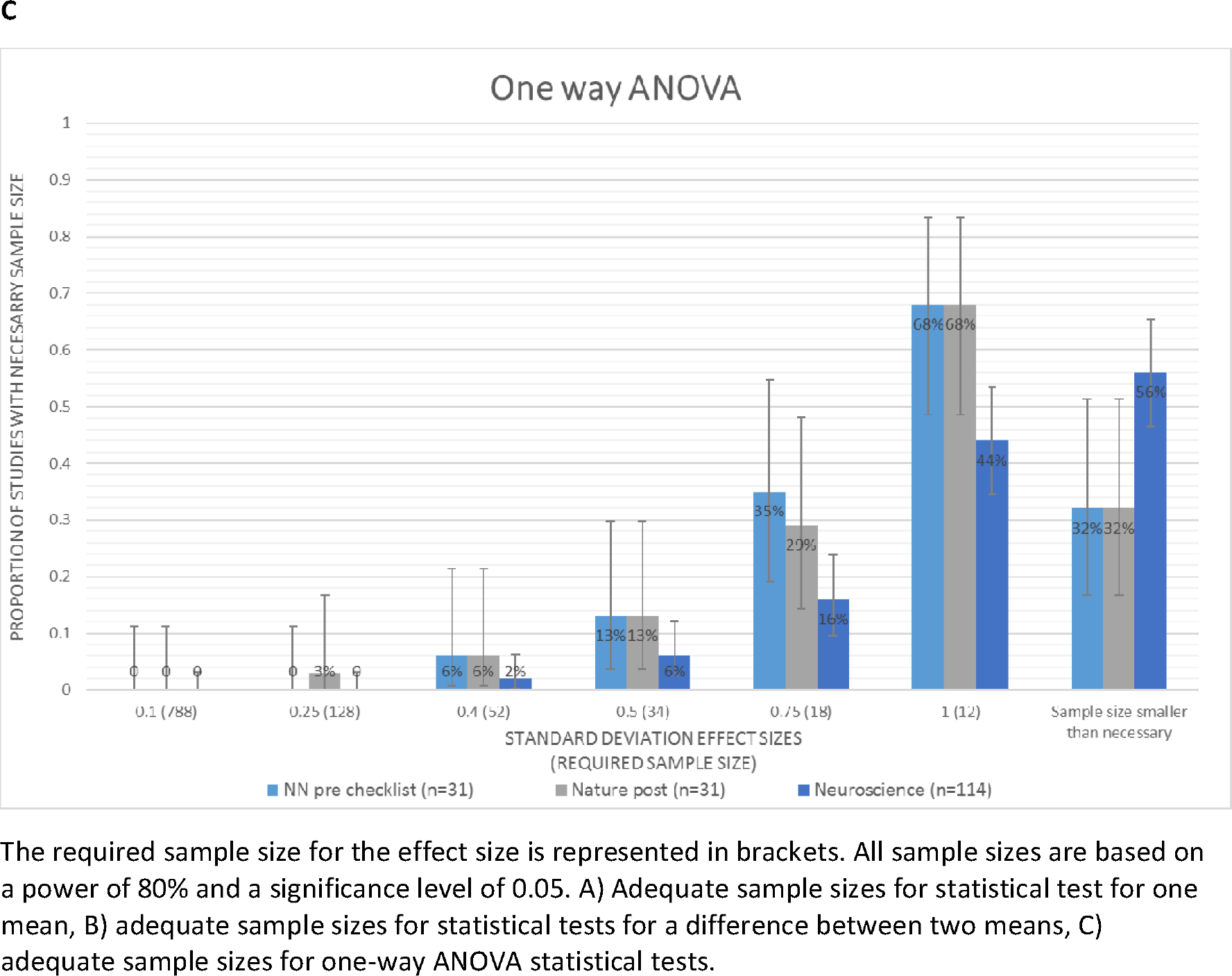
Bar chart showing the cumulative proportion of reviewed studies with adequate sample sizes for each journal.

*Neuroscience* had fewer studies with adequate sample size, according to our power calculations, than *Nature Neuroscience*. However, the trend of most studies only having adequate sample sizes to detect the largest effect estimates persisted. Almost all *Neuroscience* studies had adequate sample sizes for large effect sizes (>2 SD) when carrying out one-mean statistical tests; however, this decreased from 98% to 44% when considering two-mean or ANOVA statistical tests. This may be reflective of the actual tests carried out in the studies, or an artefact of the sample sizes presented.

## Discussion

Our analysis suggests that while the number of studies reporting a justification of their sample size in the article main text increased after checklists were implemented from 22% to 53%, the number of actual power calculations decreased from 16% to 9%. However, of the 24 studies (53%) with comments on how their sample sizes were determined, 20 made simple statements declaring no statistical methods (i.e., a formal power calculation) were used or that the sample sizes were similar to those typical for the field. This indicates that although the checklist has had the intended consequence of ensuring authors improve the transparency of approaches and methods, there has been little improvement in carrying out methods that will ultimately improve research reproducibility and reliability. When assessing whether sample sizes were adequate, the majority of studies (90-98%) were well powered to detect large effect sizes (2.0 SD) for one mean. However, few studies were adequately powered to detect smaller (and arguably more realistic) effect sizes (0.2-1.0 SD). When assessing two means, only 44-68% of studies were large enough to detect large effect sizes with 80% power. This is the same when we consider one-way ANOVAs, a common analytical method. There are no clear improvements in adequate sample sizes after checklists were implemented.

The initial *Nature Neuroscience* article introducing the checklist suggests that it is designed to promote transparency of study reporting (9). However, the results presented here indicate that an intervention intended to promote transparency has had no (or even a negative) impact on the quality of study reporting or conduct of the studies themselves. Since the number of statements on sample size increased, this suggests inadequate justification of study sample size has also increased. In our opinion, simply enforcing a checklist without checking that it both achieves the intended outcome (i.e., greater transparency and improved study design) and does not result in unintended negative consequences will not benefit the field. Over one third (38%) of the published studies still made no mention of how they determined their sample size, yet they were published alongside articles that used the checklist to better improve the transparency of their studies, even when no power calculations were carried out. If studies can be published in high profile journals without adequately considering *a priori* the sample size necessary to adequately support the planned inferential statistics, it sets a precedent that this is an appropriate approach However, if a high profile journal were to take a bold stance on this, it might encourage researchers to seek a better understanding of how and why they should implement *a priori* power calculations in to their work.

Submission checklists can be valuable tools in improving scientific quality. However, if they are treated as an end in themselves (i.e., a box-ticking exercise), rather than a means to an end (i.e., to improve the quality of reporting and more importantly to improve study design and conduct), they will not achieve this ultimate goal of improving scientific quality. Here it seems the checklist provided authors with the opportunity to give a weaker sample size justification than they otherwise might have. Our results suggest that greater transparency alone, in the absence of either improved methodological training or more rigorous enforcement by journals, may in fact take us backwards in our attempts to improve reproducibility.

Our review has only considered studies that were ultimately published in these journals, along with the detail provided in these published versions. The adequacy of sample sizes presented here is likely to be an over-optimistic estimate; we took the largest reported sample size of any one experimental group, which is not necessarily reflective of most experimental groups in the study. We also only looked at adequate sample sizes for 80% power, rather than more stringent power levels of 90% or 95%. A limited number of statistical tests were carried out for the indicative sample size calculations in this review, which may not truly reflect those carried out by each study; however, those included in this review reflect the most common tests carried out.

It is common to carry out secondary analysis of existing datasets, in which case the sample size is often fixed prior to conducting any new analyses. However, it remains possible to carry out *a priori* power calculations to identify the maximum possible power of the data given a range of plausible effect sizes. It is widely accepted that *post hoc* (or observed) power calculations should not be carried out, because observed power does not provide any additional information over the P-value alone (12). However, even given a fixed sample size, considerations of power can still be made prior to analyses being carried out to determine whether the planned study is likely to be adequately powered, or to give readers of the study a sense of the effect sizes that the study was able to detect, so that they can form their own opinion as to whether these are credible. Where studies are carrying out new data collection, power calculations should always be carried out.

Our review suggests checklists have not improved sample sizes or authors' statistical justification of their sample size. Many *Neuroscience* studies remain underpowered, resulting in the potential for many published results to be false positives, or to erroneously conclude that there is evidence of no effect when in fact there is just insufficient evidence of any effect. Journals need to consider strict enforcement of the checklists to make meaningful differences. Statistical power is a critical design consideration. Therefore, authors should be considering sample size and power at this stage of the research process, as opposed to at the publication stage, whether collecting new data or deciding whether to proceed with an analysis of existing data.

## Author Contributions

ARC carried out all reviewing of manuscripts and analysis of the data, participated in the study design and drafted the manuscript. KT and MRM conceived the study, assisted in interpreting the results and made comments on and assisted in drafting the manuscript. All authors gave final approval for publication.

